# SeuratExtend: Streamlining Single-Cell RNA-Seq Analysis Through an Integrated and Intuitive Framework

**DOI:** 10.1101/2024.08.01.606144

**Authors:** Yichao Hua, Linqian Weng, Fang Zhao, Florian Rambow

## Abstract

Single-cell RNA sequencing (scRNA-seq) has revolutionized the study of cellular heterogeneity, but the rapid expansion of analytical tools has proven to be both a blessing and a curse, presenting researchers with significant challenges. Here, we present SeuratExtend, a comprehensive R package built upon the widely adopted Seurat framework, which streamlines scRNA-seq data analysis by integrating essential tools and databases. SeuratExtend offers a user-friendly and intuitive interface for performing a wide range of analyses, including functional enrichment, trajectory inference, gene regulatory network reconstruction, and denoising. The package seamlessly integrates multiple databases, such as Gene Ontology and Reactome, and incorporates popular Python tools like scVelo, Palantir, and SCENIC through a unified R interface. SeuratExtend enhances data visualization with optimized plotting functions and carefully curated color schemes, ensuring both aesthetic appeal and scientific rigor. We demonstrate SeuratExtend’s performance through case studies investigating tumor-associated high-endothelial venules and autoinflammatory diseases, and showcase its novel applications in pathway-Level analysis and cluster annotation. SeuratExtend empowers researchers to harness the full potential of scRNA-seq data, making complex analyses accessible to a wider audience. The package, along with comprehensive documentation and tutorials, is freely available at GitHub, providing a valuable resource for the single-cell genomics community.

**Practitioner Points:**

- SeuratExtend streamlines scRNA-seq workflows by integrating R and Python tools, multiple databases (e.g., GO, Reactome), and comprehensive functional analysis capabilities within the Seurat framework, enabling efficient, multi-faceted analysis in a single environment.
- Advanced visualization features, including optimized plotting functions and professional color schemes, enhance the clarity and impact of scRNA-seq data presentation.
- A novel clustering approach using pathway enrichment score-cell matrices offers new insights into cellular heterogeneity and functional characteristics, complementing traditional gene expression-based analyses.

## INTRODUCTION

In recent years, single-cell RNA sequencing (scRNA-seq) has revolutionized our understanding of cellular diversity and complexity at an unprecedented scale across various biological disciplines. The rapid advancement of this technology has led to an explosion of computational tools and algorithms, with over 1,700 methods reported as of April 2024 (Zappia and Theis 2021). While this proliferation underscores the field’s vibrancy, it also presents a daunting challenge for researchers, who often find themselves overwhelmed by the myriad of choices and complexities.

The most common analytical tasks in scRNA-seq include doublet removal, denoising, batch integration, cell clustering and annotation, pathway and functional analysis, gene regulatory network inference, trajectory and pseudotime analysis, and cell-cell communication (Luecken and Theis 2019; Heumos et al. 2023; Kharchenko 2021; Hao et al. 2021). Each of these analytical dimensions addresses fundamental biological questions but often requires navigating a complex landscape of software tools, each with unique input requirements, operational intricacies, and output formats. Furthermore, evaluating the performance of these diverse tools across different datasets, assessing their computational resource demands, and optimizing their utilization are all critical considerations. This diversity, although scientifically enriching, frequently translates into steep learning curves and substantial barriers to entry for newcomers and even for experienced users who face interoperability issues among different tools.

Beyond the challenges of understanding and navigating these tools and algorithms, practical implementation presents its own set of hurdles. A significant obstacle in scRNA-seq analysis lies in the accessibility and user-friendliness of existing tools. Researchers often encounter issues such as convoluted code that is difficult to comprehend, non-functional scripts, challenging error-tracing, outdated software or dependencies from non-official sources that cause installation or environment conflicts, and ambiguous tutorials or function documentation, leading to significant time wastage. Moreover, mainstream scRNA-seq analysis tools are primarily developed for either the R or Python platforms, with additional options like Nextflow and Snakemake. While R, with its core package Seurat, and Python, with its central package Scanpy, have their respective strengths (Hao et al. 2021; Wolf, Angerer, and Theis 2018), proficiency in both languages is rare, posing a significant challenge, particularly for newcomers. Cross-platform tool interoperability remains a persistent issue. Additionally, the increasing adoption of Python has led to the emergence of ecosystems like scverse, centered around Scanpy (Wolf, Angerer, and Theis 2018; Virshup et al. 2023), greatly promoting the Python community’s development. In contrast, the R ecosystem surrounding Seurat appears relatively limited, even with resources like SeuratWrappers (https://github.com/satijalab/seurat-wrappers), a collection of community-provided methods and extensions curated by the Satija Lab, which offers a more limited scope compared to scverse.

To address these challenges, we present SeuratExtend, a comprehensive and integrated R package designed to streamline scRNA-seq data analysis workflows. Built upon the widely adopted Seurat framework, SeuratExtend aims to provide a user-friendly and accessible solution by focusing on essential visualization and analysis features from a practical standpoint. By leveraging the familiarity and extensive user base of Seurat, SeuratExtend offers a cohesive ecosystem that significantly enhances its capabilities. The seamless integration with Seurat and the intuitive design of SeuratExtend make it easier for R users to adopt and learn the package, thereby lowering the entry barrier for the R community.

SeuratExtend’s core philosophy emphasizes *integration*, *intuitive design*, and *visual aesthetics*. *Integration* refers to the package’s comprehensive coverage of the most practical analysis and visualization tools for scRNA-seq, incorporating essential databases like GO and Reactome (The Gene Ontology Consortium 2019; Gillespie et al. 2022), along with accompanying functions. It also seamlessly integrates classic Python tools such as scVelo, CellRank, Palantir, and SCENIC into the R environment (Bergen et al. 2020; Lange et al. 2022; Setty et al. 2019; Aibar et al. 2017), eliminating the need for proficiency in Python. *Intuitive design* and ease of use are achieved through straightforward functions complemented by detailed documentation and tutorials, enabling newcomers to effortlessly adopt the package. *Visual aesthetics*, in addition to conveying scientific information, emphasize data visualization appeal. The most common visualizations include dimension reduction plots (e.g., UMAP), violin/box plots, heatmaps, bar plots/pie charts (displaying distributions), bubble plots, waterfall plots/volcano plots, and circlize plots/river plots (showcasing interactions/connections). While Seurat, as a comprehensive tool, provides various visualization functions, there is still room for improvement. SeuratExtend significantly enhances and optimizes many of these visualizations, with a particular focus on exploring and refining color schemes. In this manuscript, we systematically present the key features and functionalities of SeuratExtend, illustrated through diverse case studies that showcase its utility in addressing prevailing challenges and enhancing the scRNA-seq data analysis experience across various applications.

## MATERIAL AND METHODS

### Integration of multiple databases for GSEA

The Gene Ontology (GO) database was obtained from the official website (https://geneontology.org/), including the .gaf and .obo files for both human and mouse. The .gaf files contain pathway-gene information, while the .obo files contain GO term naming, definitions, and hierarchical relationships between terms. The msga package’s readGAF function and ontologyIndex’s get_OBO were used to process the respective file formats. To construct the hierarchy, obsolete GO terms not present in the relational network were removed, and ontologyIndex was used to build standardized ontology_index objects for subsequent analyses.

For the Reactome database, relevant files were downloaded from the official website (https://reactome.org/), including “Ensembl2Reactome_PE_All_Levels.txt” (containing pathway-gene information) and “Ensembl2Reactome_PE_Reactions.txt” (containing information on relationships between pathways). Human and mouse-related pathways were extracted separately. As gene names were in Ensembl ID format, they were converted to gene symbols, and ontologyIndex was used to construct standardized ontology_index objects for subsequent analyses.

Other databases, such as Hallmark 50, KEGG, and BioCarta, were downloaded from https://www.gsea-msigdb.org/ and standardized into a uniform format. Cell type-marker gene information was obtained from the PanglaoDB database, downloaded from https://panglaodb.se/.

### Seurat object integration and conversion

The hdf5r package was used to create a dataset named “LOOM_SPEC_VERSION” in an HDF5 file, setting its value to “3.0.0” to specify the version of the Loom file format being used. The data type for this dataset was set to UTF-8 string. Count matrices (raw, normalized, or velocyto’s spliced/unspliced) were stored in the layers group, while meta.data information was stored in the col_attrs group. To convert from Loom to AnnData, Python’s Scanpy read_loom function was used.

### Python environment setup and Integration of Python tools using conda and reticulate

The reticulate framework was used to integrate Python in R. By default, reticulate was used to create a conda environment named “seuratextend,” containing all potentially required packages. The conda environment was created and tested on Linux, macOS, and Windows systems to ensure compatibility and avoid version conflicts between Python packages. The environment specifications were saved as .yml files to allow conda to install an identical environment on the user’s respective operating system.

Visualizations for scVelo and CellRank were generated using reticulate to call the respective packages. For Palantir and MAGIC, reticulate was used to perform the computations, and the results were exported using numpy and imported into the Seurat object in R. SCENIC’s loom output was imported into the Seurat object using the LoomR package, creating an assay named “TF.” Various SeuratExtend visualization functions were then used to generate plots. For gene regulatory network visualizations, such as Figure 3G, CytoScape was used in conjunction with SeuratExtend.

### Enhanced visualization

Visualization tools such as heatmaps, dimensional reduction plots, violin/box plots, cluster distribution plots, waterfall plots, and GSEA plots were built using the ggplot2 package, with layouts adjusted using cowplot. All statistical calculations were performed using the ggpubr package.

### Implementation of professional color schemes

The “color_pro” presets were constructed using the I Want Hue tool (http://medialab.github.io/iwanthue/). The provided API and JavaScript code were used to generate all presets. The specific code is available in Supplementary Data 1.

### Gene identifier conversion

Gene naming conversions between human and mouse gene symbols and Ensembl IDs were performed using the BioMart database. The biomaRt package was utilized for conversions, with improvements in reliability and performance achieved by localizing the most commonly used databases, eliminating the need for internet connectivity and addressing frequent instability issues with biomaRt. For UniProt ID conversion, the database was downloaded from the official website (https://www.uniprot.org/) and used for conversion to gene symbols.

## RESULTS

### SeuratExtend: A Comprehensive R Ecosystem for Single-Cell Analysis

Building upon the foundation laid by Seurat, SeuratExtend aims to create a more efficient and integrated workflow for single-cell RNA sequencing analysis in the R environment. While the Python ecosystem has benefited greatly from the comprehensive scverse project (Virshup et al. 2023), which utilizes the universal AnnData format to seamlessly connect various tools and algorithms, a comparable integrated solution has been lacking in the R community. SeuratExtend addresses this gap by providing a unified framework centered around the Seurat object, effectively becoming the R counterpart to scverse.

SeuratExtend expands upon the Seurat framework by integrating a comprehensive suite of tools and functionalities (**Figure 1**). The package encompasses four key areas: (1) an advanced functional and pathway analysis module, which incorporates multiple databases and utilizes the AUCell algorithm for gene set enrichment analysis; (2) seamless integration of Python-based tools, enabling sophisticated analyses such as trajectory inference, gene regulatory network reconstruction, and data denoising; (3) enhanced visualization capabilities, featuring optimized plotting methods and professionally curated color schemes; and (4) a collection of utility functions that streamline common tasks in single-cell analysis workflows. This integrated approach not only simplifies complex analytical processes but also bridges the gap between R and Python environments, providing researchers with a versatile toolkit for comprehensive single-cell RNA-seq data analysis.

**Figure 1.**
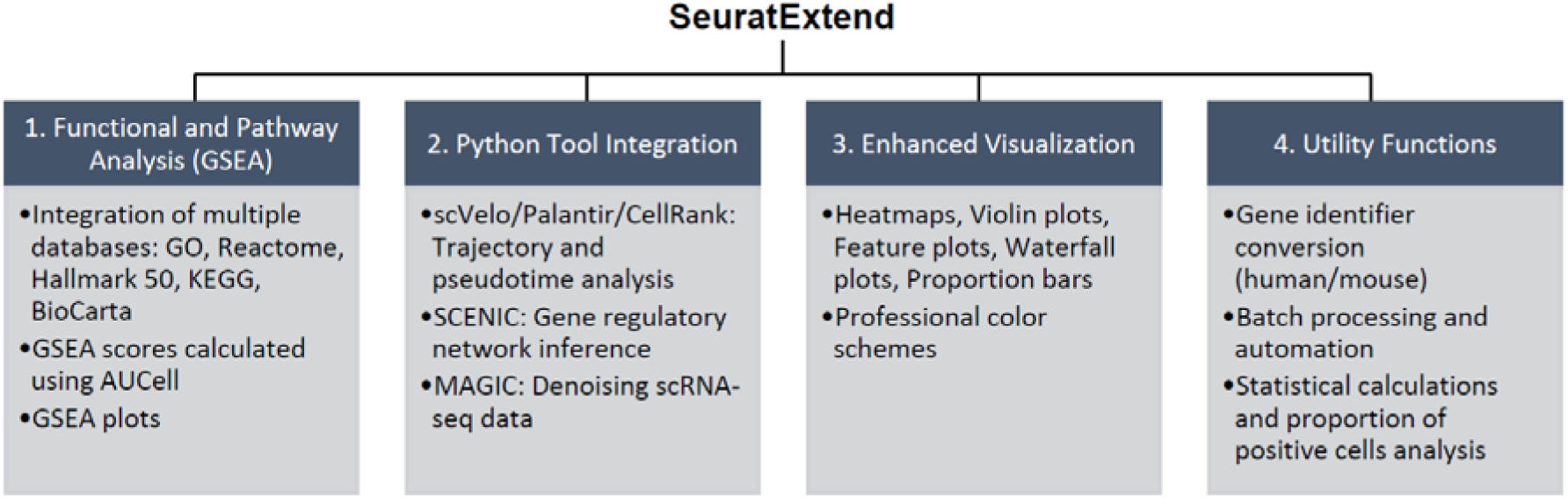
Overview of the SeuratExtend package’s key features. SeuratExtend streamlines single-cell RNA-seq data analysis by integrating essential components into the Seurat framework: (1) Functional and Pathway Analysis (GSEA) with multiple databases and AUCell algorithm; (2) Python Tool Integration for trajectory analysis (scVelo, Palantir, CellRank), gene regulatory network inference (SCENIC), and denoising (MAGIC); (3) Enhanced Visualization with optimized methods and professional color schemes; and (4) Utility Functions for gene identifier conversion, batch processing, and statistical analysis.

To illustrate the unique position of SeuratExtend in the single-cell analysis toolkit landscape, we conducted a comprehensive comparison with other prominent tools and frameworks (**Table 1**). Compared to Seurat and SeuratWrappers, it offers a higher level of integration, providing a more seamless workflow from data input to final analysis and visualization. While other platforms like Snakemake offer powerful workflow management capabilities, they often require learning custom languages, presenting a steep learning curve for many researchers. SeuratExtend, in contrast, employs the widely-used R language, making it more accessible to a broader range of scientists, particularly those with backgrounds in biology and statistics. Furthermore, SeuratExtend uniquely bridges the gap between R and Python ecosystems. Unlike scverse, which primarily focuses on Python-based tools, SeuratExtend provides direct functions for converting between Seurat objects and AnnData format, facilitating interoperability between R and Python environments. This feature allows users to leverage powerful Python-based tools directly within the R environment, eliminating the need for researchers to learn Python or switch between different platforms. Consequently, this high level of integration not only expands the toolkit available to R users but also streamlines the workflow by enabling a more cohesive analytical process, making SeuratExtend a versatile and user-friendly option in the diverse landscape of bioinformatics tools.

**Table 1.** Comparison of SeuratExtend with other single-cell analysis toolkits.

| Features | SeuratExtend | scverse | Seurat/<br>SeuratWrappers | scRNA-tools | Snakemake |
| --- | --- | --- | --- | --- | --- |
| **Primary Language** | R | Python | R | Various | Custom |
| **Unified Data Structure** | √ (Seurat) | √ (AnnData) | √ (Seurat) | × | × |
| **R-Python Integration** | √ | Limited | Limited | × | Varies |
| **Comprehensive GSEA Integration** | √ | Limited | Limited | Varies | Varies |
| **Enhanced Visualization** | √ (ggplot) | Varies | Limited | Varies | Varies |
| **Learning Curve** | Moderate | Moderate | Moderate | Varies | Steep |
| **Integrated Tutorial** | √ | √ | √ | Varies | Varies |

It’s important to emphasize that SeuratExtend is not merely a collection of existing tools. It introduces numerous extensions and novel developments, including enhanced database integration, improved visualization capabilities, and a suite of practical utility functions. These innovations, combined with its integrative approach, position SeuratExtend as a powerful and unique resource in the R ecosystem for single-cell analysis. In the following sections, we will delve deeper into the key components and functionalities of SeuratExtend, demonstrating how it streamlines and enhances the single-cell RNA-seq analysis workflow.

### Integrated Functional Enrichment Analysis with Multi-Source Databases

SeuratExtend offers an integrated approach to functional enrichment analysis, incorporating multiple authoritative databases within a unified framework (**Figure 1**). This feature enables researchers to harness diverse data sources and employ state-of-the-art analytical methods, facilitating a comprehensive understanding of their single-cell transcriptomic data.

At the core of this integration lies the Gene Ontology (GO) and Reactome databases. The GO database provides a structured representation of biological processes, molecular functions, and cellular components, enabling researchers to interpret their gene expression data within the context of well-established biological knowledge (The Gene Ontology Consortium 2019). Complementing the GO database, SeuratExtend also integrates the Reactome knowledgebase, a comprehensive resource for curated biological pathways (Gillespie et al. 2022). This integration allows researchers to explore the functional implications of their data within the context of well-described cellular processes, signaling cascades, and disease mechanisms. Furthermore, SeuratExtend incorporates a range of additional databases, including the Hallmark 50, KEGG, and BioCarta, providing researchers with an extensive collection of curated gene sets for their analyses.

Traditionally, functional enrichment analysis for bulk RNA-seq data involves identifying differentially expressed genes (DEGs) based on a predetermined cut-off and then comparing the resulting gene list against pathway databases to calculate enrichment scores. However, this approach suffers from inherent biases and limitations, as it fails to capture the nuances of single-cell data and relies on arbitrary cut-offs. SeuratExtend addresses this challenge by employing a dedicated single-cell-based method, AUCell, specifically designed to exploit the unique characteristics of single-cell RNA-sequencing data. Unlike bulk-sample-based methods, which can lead to biases, single-cell-based methods like AUCell are less susceptible to such issues. A comprehensive benchmarking study (Noureen et al. 2022) evaluated various supervised signature-scoring methods, including ssGSEA, AUCell, Single Cell Signature Explorer (SCSE), and Jointly Assessing Signature Mean and Inferring Enrichment (JASMINE). The study concluded that while bulk-sample-based methods can introduce biases when applied to single-cell RNA-sequencing data, single-cell-based methods, such as AUCell and JASMINE, exhibit superior performance. Considering its adaptability, compatibility with downstream analyses (e.g., SCENIC), and widespread recognition within the scientific community, SeuratExtend employs AUCell as its primary algorithm for calculating gene set enrichment scores.

The integrated functional enrichment analysis in SeuratExtend is designed to be accessible and user-friendly. Researchers can easily navigate and search through the available databases, filter pathways based on specific criteria (e.g., gene count, parent terms, or end-level terms). SeuratExtend constructs comprehensive hierarchical structures for the GO and Reactome databases, enabling researchers to perform targeted analyses on specific categories of interest, such as immune processes, metabolism, or signal transduction pathways, rather than analyzing the entire database in a non-specific and computationally intensive manner. Researchers can perform GSEA on these specific categories or any other term from the GO hierarchy, and visualize the results through informative heatmaps, violin plots, waterfall plots, or plots that emulate the GSEA plot developed by the Broad Institute (**Figures 2A-2C**). Additionally, SeuratExtend provides complementary functions to convert cryptic pathway identifiers into more interpretable names, further enhancing the user experience. This integration enables researchers to tap into the collective knowledge from multiple authoritative sources or custom gene sets, facilitating a deeper understanding of the biological processes underlying their single-cell transcriptomic data.

**Figure 2.**
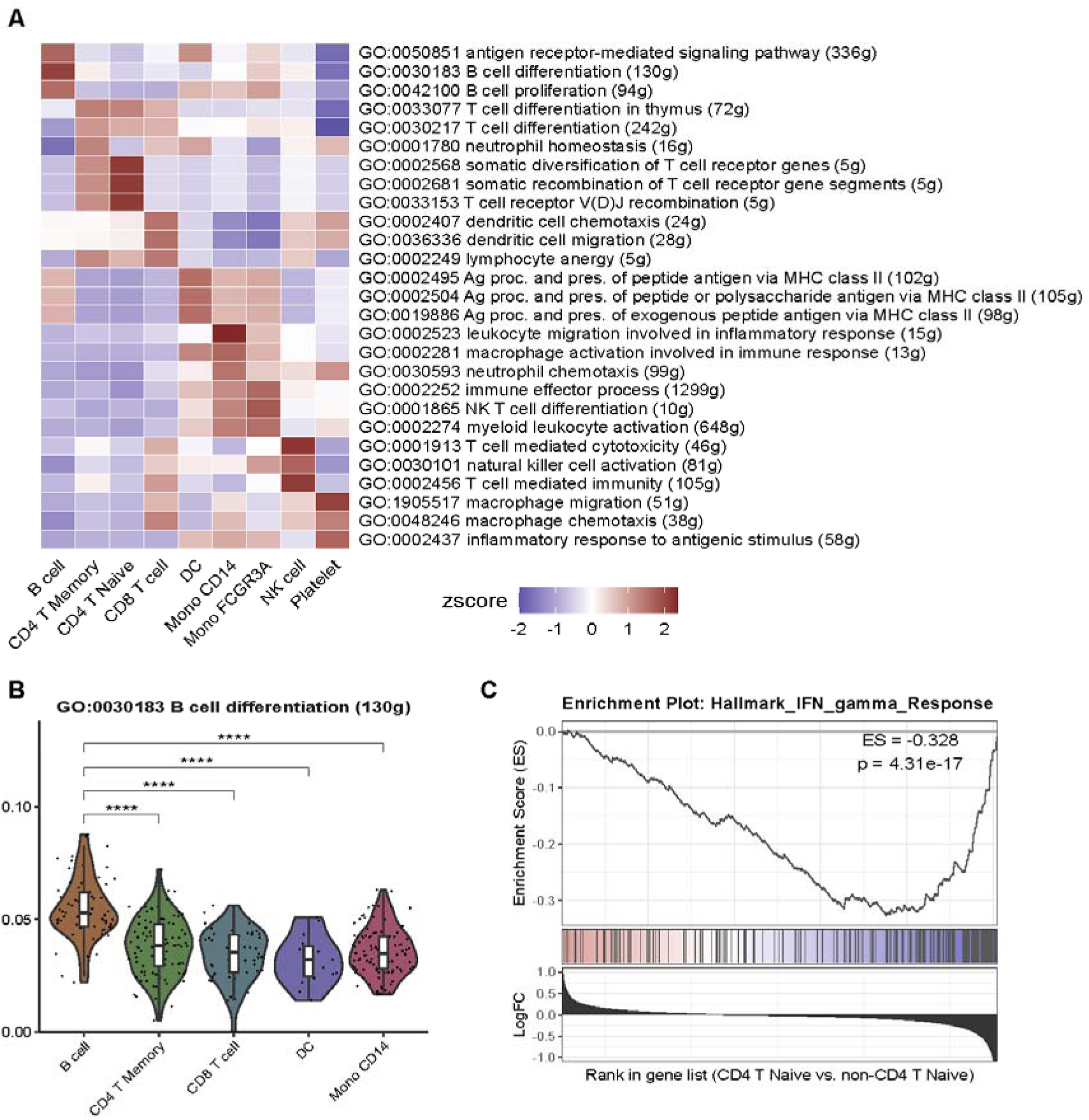
Visualizations of gene set enrichment analysis (GSEA) results using SeuratExtend. (A) Heatmap displaying the z-scores of immune-related GO terms across different cell types or conditions. (B) Violin plots showing the distribution of GSEA scores for the selected GO term across different cell types, with asterisks indicating statistically significant differences (**** p < 0.001). (C) GSEA plot for the Hallmark IFN gamma Response gene set, comparing CD4 T naive cells to non-CD4 T naive cells. Kolmogorov-Smirnov Test.

### Seamless Integration of Python Tools within an R Environment

SeuratExtend offers seamless integration of Python tools within the R environment, enabling researchers to utilize the analytical capabilities of both ecosystems. This integration is particularly valuable for tasks such as trajectory analysis, pseudotime calculation, gene regulatory network inference, and denoising, where Python tools like scVelo, Palantir, CellRank, SCENIC, and MAGIC excel (Bergen et al. 2020; Setty et al. 2019; Lange et al. 2022; Aibar et al. 2017; van Dijk et al. 2018) (**Figure 1**).

To facilitate this integration, SeuratExtend implements three key components: First, it redesigns the architecture for converting between fundamental scRNA-seq data storage formats, including Seurat, loom, and AnnData objects. Second, it provides a one-stop solution for Python environment setup across Windows, macOS, and Linux platforms, freeing users from resolving environment conflicts between different tools. Third, it employs the reticulate framework to enable the execution of Python tools within R, eliminating the need for users to write Python code directly.

The Seurat object serves as the primary data storage format in R, while the AnnData object is the standard format for Python’s Scanpy library. The loom file, a widely adopted hierarchical data format for storing and exchanging large, sparse matrices and associated metadata, acts as a bridge for communication between R and Python. In earlier versions of SeuratExtend, the conversion between Seurat objects and loom files relied on packages such as SeuratDisk and LoomR. However, as these were non-CRAN packages with limited maintenance and potential instability issues, SeuratExtend gradually reduced its dependence on external sources. Instead, it utilizes the essential CRAN-sourced hdf5r package to construct functions for converting between Seurat and loom formats at a lower level, ensuring more stable operations. Similarly, the conversion from Seurat/loom to AnnData directly employs the reticulate framework and Scanpy, avoiding the use of external R packages.

Many Python tools are powerful, but resolving environment conflicts between different tools can be a significant challenge. Sometimes, users must create separate virtual or conda environments for each tool, which can be inconvenient and wasteful of disk space. Addressing these issues can be challenging even for experienced Python users, let alone those who primarily work with R. To streamline the installation and management of Python dependencies, SeuratExtend leverages the power of conda environments through the reticulate package. Users can easily install a dedicated conda environment named “seuratextend,” which automatically sets up all the required Python packages, ensuring a hassle-free integration experience across Windows, macOS, and Linux platforms.

With these components in place, SeuratExtend unlocks a wide range of analytical possibilities. Users can employ the powerful trajectory analysis capabilities of scVelo and Palantir (**Figures 3A-3D**), which utilize RNA velocity and diffusion maps to predict cellular differentiation pathways and calculate pseudotime, respectively. These are highly regarded and popular tools for trajectory and pseudotime analysis. Additionally, CellRank offers an alternative approach to trajectory analysis, utilizing pre-calculated pseudotime to create informative trajectory plots. Furthermore, due to the sparse nature of scRNA-seq data, denoising techniques play a valuable role. Studies have compared various denoising algorithms (Galuzzi, Vanoni, and Damiani 2022) and identified MAGIC (Markov Affinity-based Graph Imputation of Cells) as a promising choice for improving clustering results (**Figure 3E**). While the Rmagic package was previously available on CRAN but has since been removed due to inactivity, SeuratExtend integrates the use of MAGIC without relying on Rmagic.

**Figure 3.**
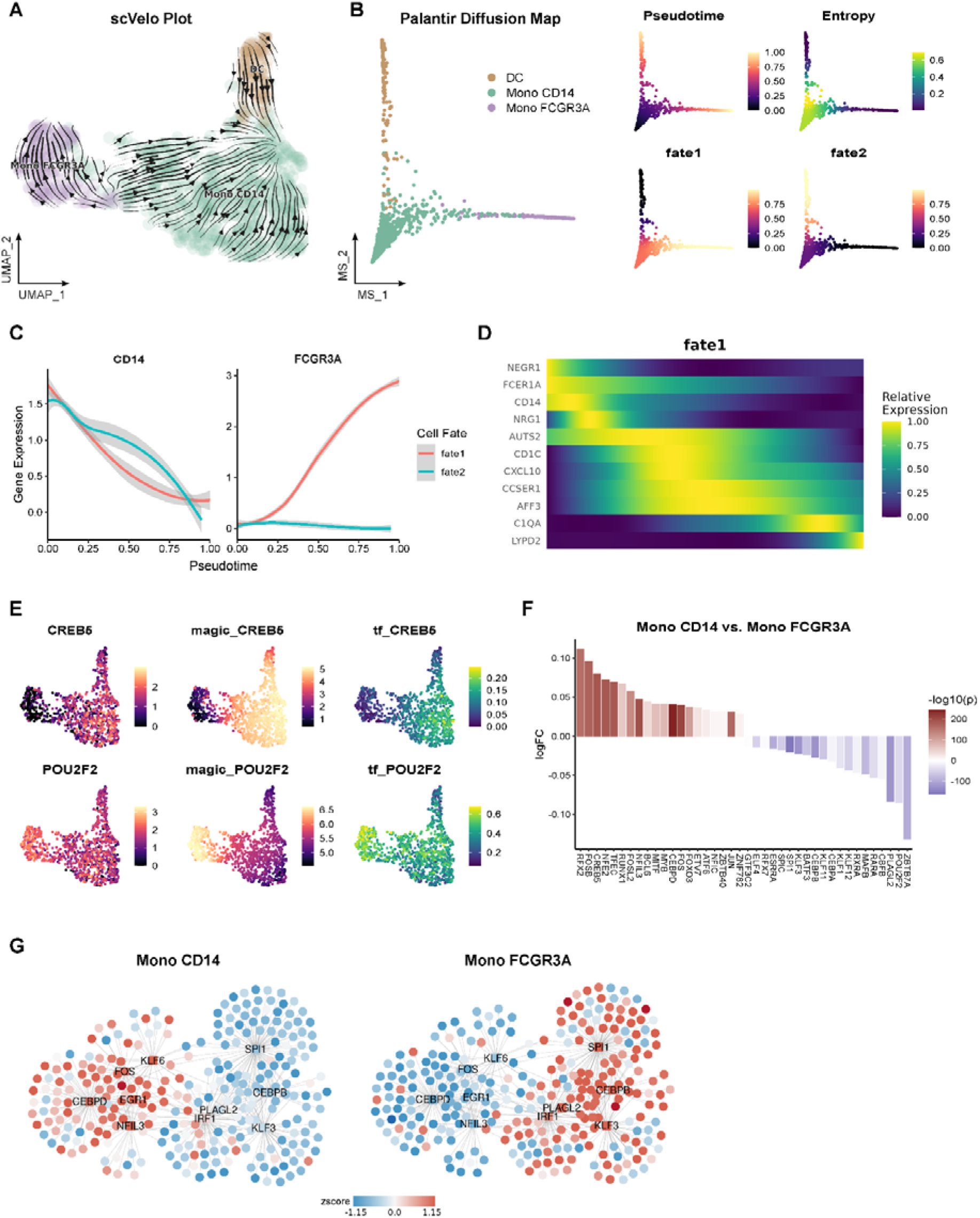
Trajectory analysis, pseudotime inference, denoising, and gene regulatory network visualization using SeuratExtend’s integrated Python tools. (A) RNA velocity analysis using scVelo, displaying velocity vectors on a UMAP embedding. (B) Diffusion map and pseudotime calculation using Palantir, comparing DC, Mono CD14, and Mono FCGR3A cell populations. (C) Gene expression dynamics along the pseudotime trajectory for the CD14 and FCGR3A genes. (D) Heatmap showing the relative expression of fate marker genes (rows) along the pseudotime trajectory (columns) for fate1. (E) UMAP visualizations of CREB5 and POU2F2 expression before (CREB5, POU2F2) and after (magic_CREB5, magic_POU2F2) denoising with MAGIC, as well as the transcription factor activity AUCell score (tf_CREB5, tf_POU2F2) inferred by SCENIC. (F) Waterfall plot highlighting differential TF regulon activities between Mono CD14 and Mono FCGR3A cell populations, with the top 20 TFs labeled. (G) Gene regulatory networks predicted by SCENIC for Mono CD14 and Mono FCGR3A cell populations, with nodes colored by relative gene expression (round nodes) or regulon activity (square nodes).

Moreover, SeuratExtend integrates SCENIC, a comprehensive computational method for inferring gene regulatory networks from single-cell transcriptomic data. SeuratExtend can directly import pySCENIC-generated loom files into Seurat objects, enabling researchers to gain insights into the intricate regulatory mechanisms governing gene expression at the single-cell level (**Figures 3E-3G**).

This seamless integration of Python tools within the R environment allows researchers to harness the collective strengths of both ecosystems, enabling them to tackle complex analytical challenges and unravel the intricate mechanisms governing cellular processes at the single-cell level.

### Enhanced Data Visualization with Aesthetic Refinement

Effective visualization is paramount in the field of single-cell transcriptomics, enabling the communication of intricate cellular patterns, facilitating data interpretation, and conveying scientific insights. SeuratExtend recognizes the importance of aesthetics in scientific communication and introduces an array of tools to enhance the visual appeal and clarity of data representations.

While offering advanced analytical capabilities, SeuratExtend also places great emphasis on optimizing and expanding the fundamental visualization functions essential for presenting complex analyses. The package provides improved versions of core visualization tools such as heatmaps, dimensional reduction plots, violin/box plots, cluster distribution plots, and waterfall plots (**Figures 1, 2A, 2B, 3E, and 3F**). These enhancements streamline the visualization process and introduce new features and customization options, such as convenient statistical annotation and layout settings, enabling researchers to create more informative and visually compelling representations of their data.

Central to SeuratExtend’s approach to visualization is the implementation of thoughtfully curated color schemes that adhere to principles of effective data science plotting. The creation of the “professional discrete color” (*color_pro*) series stems from the understanding that choosing the right colors for scientific visualizations is critical. Colors must be distinct enough to differentiate data points clearly, yet coordinated and subdued enough to maintain professionalism and avoid visual strain.

In the realm of data science visualization, certain color choices should be avoided, such as monochromatic schemes that can reduce visual distinction (**Figure 4A**), causing data points to blend together. Similarly, overly saturated colors can be visually aggressive and distracting, detracting from the scientific message (**Figure 4A**). While certain vibrant schemes might be engaging in an advertising context, they may be considered informal for professional journal standards (**Figure 4B**).

**Figure 4.**
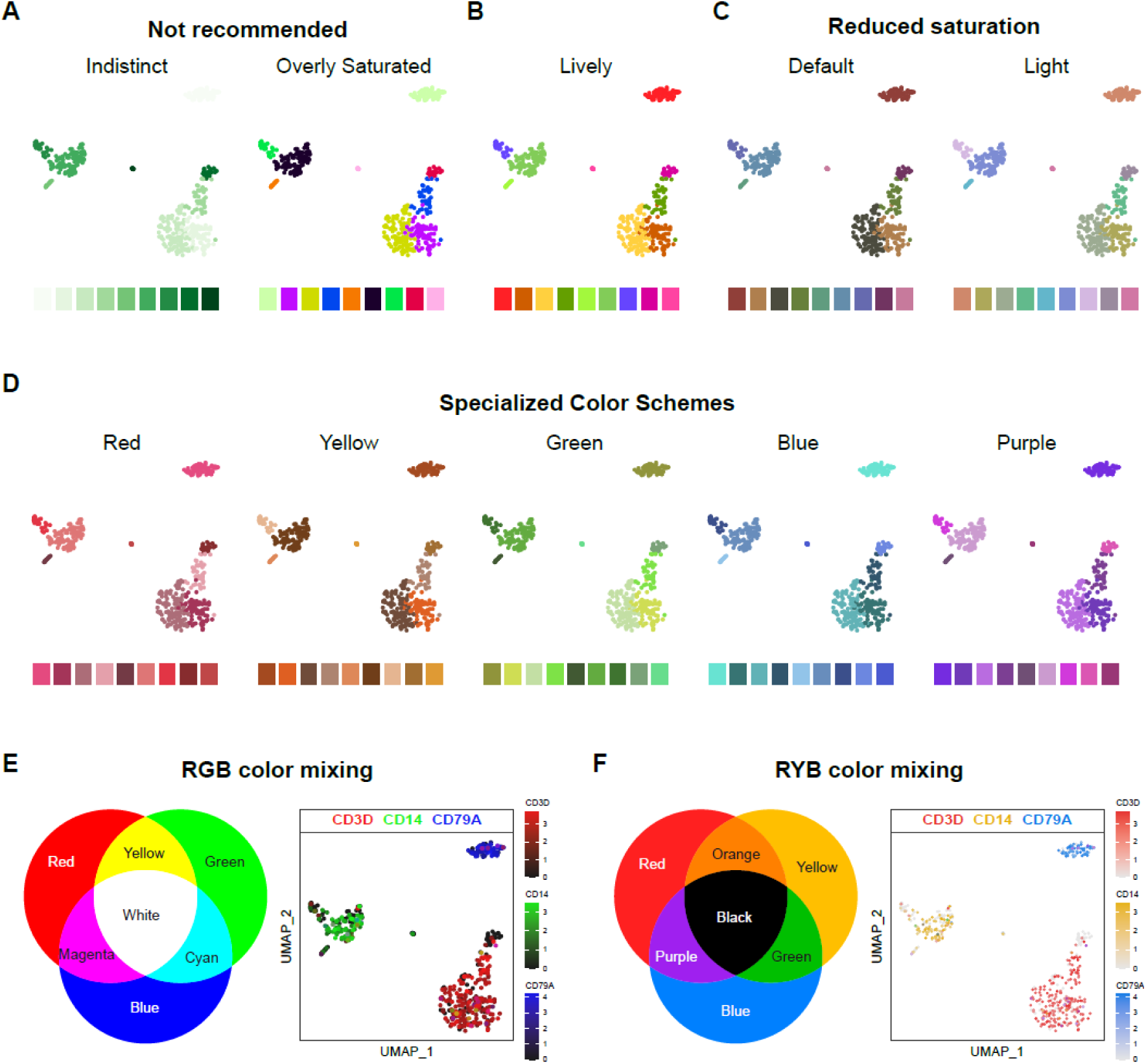
Comparison of color schemes and their suitability for data science visualization. (A) Not recommended color schemes: indistinct monochromatic colors that reduce visual distinction, and overly saturated colors that can be visually aggressive and distracting. (B) A lively color scheme that, while engaging, may be considered informal for professional journal standards. (C) The “color_pro” series offers two meticulously crafted color schemes: “default” and “light”, which span the entire hue domain and cater to general use cases. (D) The “color_pro” series also includes five specialized color schemes: “red”, “yellow”, “green”, “blue”, and “purple”, which offer harmonious hues confined to specific regions, enabling vibrant yet coordinated visualizations that reflect the biological or categorical properties of the data. (E-F) Multi-feature visualization using color mixing principles in the RGB (E) or RYB (F) color system. The expression levels of three genes (CD3D, CD14, and CD79A) are displayed on a UMAP plot.

The *color_pro* series addresses these concerns by offering a collection of seven meticulously crafted color schemes: *default*, *light*, *red*, *yellow*, *green*, *blue*, and *purple* (**Figures 4C** and **4D**). These palettes are generated using *I Want Hue* (http://medialab.github.io/iwanthue/), an optimized algorithm that ensures visually pleasing and distinctly separable color combinations, with carefully adjusted parameters to align with the aforementioned principles. The *default* and *light* schemes span the entire hue domain, catering to general use cases while accommodating different background and text color requirements (**Figure 4C**). The specialized color schemes (*red*, *yellow*, *green*, *blue*, and *purple*) offer harmonious hues confined to specific regions (**Figure 4D**), enabling vibrant yet coordinated visualizations that reflect the biological or categorical properties of the data.

In addition to the curated color schemes, SeuratExtend introduces an innovative approach to visualize multiple features simultaneously on a single dimension reduction plot, such as UMAP. This novel method utilizes the principles of color mixing in either the RGB (red, green, blue) or RYB (red, yellow, blue) color systems to represent the levels of different features, such as gene expression, pathway AUCell scores, or transcription factor activities.

In the RGB system, black represents no or low expression, while brighter colors indicate higher levels (**Figure 4E**). This color mixing approach is straightforward and well-established in the digital color space. In contrast, the RYB system, which is more intuitive and closely mimics the subtractive color mixing used in traditional art, employs white to represent no expression, with deeper colors indicating higher expression levels (**Figure 4F**).

However, designing an algorithm for RYB color mixing that accurately simulates real-world color blending poses a challenge. Moreover, the pure primary colors in the RYB system have inherent limitations for data visualization purposes, as yellow tends to be too light and blue too dark. To address these issues, SeuratExtend has developed a custom algorithm for RYB color mixing and has fine-tuned the brightness and saturation of the primary colors to optimize them for data visualization (**Figure 4F**).

In summary, SeuratExtend equips researchers to create visually compelling and scientifically rigorous representations of complex biological insights by seamlessly integrating innovative color schemes, multi-feature visualization capabilities, and intuitive color mixing principles into its optimized plotting tools.

### Utility Toolset: Streamlining Analysis Workflows

SeuratExtend introduces a comprehensive suite of utility tools designed to streamline and enhance scRNA-seq data analysis workflows (**Figure 1**). A recurring challenge in scRNA-seq analysis involves reconciling gene identifiers across disparate databases and organisms. SeuratExtend addresses this by providing robust functions that facilitate gene naming conversions between human and mouse gene symbols, Ensembl IDs, and UniProt accession numbers. These functions utilize localized databases, ensuring reliable and efficient conversion while mitigating instability issues associated with online resources. Users can directly convert gene expression matrices between human and mouse counterparts, streamlining cross-species analyses. For scenarios requiring online resources, SeuratExtend provides flexibility to fetch results directly from BioMart databases.

SeuratExtend introduces convenient tools for computing statistics and assessing the proportion of positive cells within clusters or groups. One tool computes various metrics, including mean, median, z-scores, or log-fold changes, for genomic data stored in Seurat objects or standard matrices. This enables researchers to identify cluster-specific gene expression patterns or pathway activities, crucial for understanding cellular heterogeneity and functional characteristics. Another tool allows users to assess the proportion of positive cells expressing a particular feature within specified clusters or groups, enabling the identification of genes or pathways exhibiting significant expression levels within subpopulations of cells.

To further enhance accessibility and reproducibility, SeuratExtend introduces a function that automates the execution of a standard Seurat pipeline, including normalization, principal component analysis (PCA), clustering, and uniform manifold approximation and projection (UMAP) visualization. This function offers extensive customization options and intelligent conditional execution, ensuring that specific steps are re-run only when necessary. Additionally, it provides the option to integrate and correct for batch effects using the Harmony algorithm. According to benchmarking studies (Yu et al. 2023), Harmony performs well in general tasks, offering fast computation and efficient resource utilization while maintaining simplicity of use, making it an excellent choice for batch effect correction in scRNA-seq data analysis.

SeuratExtend’s utility toolset represents a valuable advancement in streamlining and enhancing scRNA-seq data analysis workflows, facilitating efficient navigation of the complexities of scRNA-seq data exploration and accelerating scientific discoveries.

### Comprehensive Tutorials and Interactive Learning Resources

SeuratExtend prioritizes accessibility and user-friendliness, reflected in its comprehensive tutorials and interactive learning resources. A meticulously crafted, in-depth tutorial (https://huayc09.github.io/SeuratExtend/) serves as a comprehensive guide, catering to both novice and experienced users. Beyond its utility for SeuratExtend, the tutorial doubles as a valuable educational tool for scRNA-seq analysis in general, combining theoretical concepts with practical implementations.

To further enhance the user experience, SeuratExtend incorporates large language models (LLMs). By training an LLM on the package’s documentation and tutorials, the developers have created an interactive chatbot (https://chatgpt.com/g/g-8scQjmzkd-scrna-seq-assistant) capable of answering users’ queries and providing guidance. This virtual assistant offers real-time support, addressing questions and deepening users’ understanding of SeuratExtend’s features, ultimately enhancing productivity and accelerating research endeavors.

### Evolving Applications of SeuratExtend: From Early Adoption to Current Capabilities

SeuratExtend’s development has been driven by real-world research needs, with its capabilities expanding and refining over time. Here, we present two case studies that demonstrate the application of early versions of SeuratExtend in addressing complex biological questions across diverse research domains. These studies not only showcase the package’s initial utility but also serve as a foundation for understanding its subsequent evolution and enhancement.

In a study by Hua-Vella et al. (2022), an early version of SeuratExtend was employed to investigate the transcriptional landscape of tumor-associated high-endothelial venules (TU-HEVs), specialized vascular structures facilitating lymphocyte infiltration into tumors (Hua et al. 2022). The study integrated multiple single-cell RNA-sequencing datasets from murine and human tumor endothelial cells (ECs). SeuratExtend facilitated the identification of distinct EC subtypes, including TU-HEVs, lymph node HEVs (LN-HEVs), and tumor ECs (TU-ECs). Through differential expression and gene set enrichment analyses, unique transcriptional signatures associated with each subtype were revealed, shedding light on the specialized functions of TU-HEVs in lymphocyte recruitment and antigen presentation. The package’s trajectory inference capabilities were instrumental in uncovering that TU-HEVs arise from the metaplastic conversion of postcapillary venules (PCVs) in response to immunotherapy-induced signals. This finding, corroborated by in vivo lineage tracing experiments, highlighted the dynamic nature of TU-HEV formation. Furthermore, SeuratExtend enabled the exploration of TU-HEV-associated lymphocyte niches, revealing their role in fostering the expansion and differentiation of progenitor CD8+ T cells (pTEX) into cytotoxic effector T cells (tTEX). This analysis integrated transcriptomic data with advanced imaging techniques, demonstrating SeuratExtend’s flexibility in handling diverse data types.

In a subsequent study by Hua et al. (2023), SeuratExtend was utilized to decipher complex immune responses in patients with rare systemic autoinflammatory diseases (SAIDs) undergoing anti-TNF therapy (Hua et al. 2023). The package facilitated detailed transcriptomic profiling of peripheral blood mononuclear cells (PBMCs) and polymorphonuclear neutrophils (PMNs). Leveraging SeuratExtend’s analytical tools, the researchers identified unexpected increases in TNF and IL-1 levels post-therapy, challenging conventional understanding of disease activity markers. This finding prompted further investigations into alternative therapeutic response mechanisms. The package’s trajectory analysis capabilities were crucial in exploring macrophage differentiation pathways. This analysis led to the proposal of a novel therapeutic response mechanism, suggesting that anti-TNF therapy inhibits macrophage differentiation rather than solely targeting cytokine suppression.

These case studies, along with feedback from many researchers who have tested the package, have underscored SeuratExtend’s practical value in single-cell RNA-seq analysis. Recognizing its potential to benefit a wider scientific community, we have committed to continuously improving and expanding its capabilities. Since these initial applications, SeuratExtend has undergone significant development. Many functions have been restructured to enhance code maintainability and optimize computational efficiency. Compatibility with different versions of R and Seurat has been improved, ensuring broader usability across various research environments. Moreover, we have refined existing functions and introduced new ones to address emerging analytical needs in the rapidly evolving field of single-cell genomics.

A key focus of recent development has been on improving user accessibility. While early versions of SeuratExtend provided a set of useful functions, they lacked comprehensive documentation and tutorials, which limited their widespread adoption. To address this, we have created extensive documentation and step-by-step tutorials, significantly enhancing the package’s usability and lowering the entry barrier for researchers new to single-cell analysis.

The current version of SeuratExtend represents a substantial advancement over its earlier iterations, offering a more comprehensive, integrated, and user-friendly platform for single-cell RNA-seq analysis. As we continue to refine and expand SeuratExtend, our goal is to provide the scientific community with an increasingly powerful and accessible tool for uncovering biological insights from complex single-cell datasets. This ongoing development ensures that SeuratExtend remains at the forefront of single-cell analysis methodologies, adapting to the evolving needs of researchers in this dynamic field.

### Novel Applications of SeuratExtend in Pathway-Level Analysis and Cluster Annotation

The SeuratExtend framework offers innovative approaches to address fundamental challenges in single-cell transcriptomic data analysis, such as cellular heterogeneity interpretation and cluster annotation. By leveraging the package’s comprehensive integration of pathway databases and analytical tools, researchers can explore novel strategies to gain deeper insights into the functional characteristics of cell populations and streamline the annotation process.

### Exploring and Analyzing Single-Cell Data at the Pathway Level

SeuratExtend introduces a novel approach to dimensionality reduction and clustering by harnessing its extensive integration of pathway databases, such as Gene Ontology (GO), Reactome, KEGG, and BioCarta. Instead of relying solely on gene expression-cell matrices, this method transforms the data into pathway enrichment score-cell matrices using AUCell. Subsequent analyses, including PCA and clustering, are then performed on these pathway-level matrices. This pathway-based approach offers several advantages. First, clustering based on pathway-level information rather than individual genes greatly reduces the impact of gene expression fluctuations. Second, by focusing on curated pathways containing functionally characterized genes, the influence of sample-specific genes with unknown functions, such as non-coding RNAs and pseudogenes, is minimized.

To illustrate the potential of this approach, we present an example using scRNA-seq data from a melanoma cohort (Pozniak et al. 2024). When clustering is based on gene expression, the UMAP visualization is significantly influenced by sample origin (**Figure 5A**). In contrast, clustering based on pathway enrichment scores derived from multiple databases (GO, Reactome, KEGG, and BioCarta) effectively aggregates cells with conserved features, such as T cells and B cells, while preserving the biological variations in malignant cells across samples (**Figure 5B**). Importantly, this pathway-based analysis can provide novel functional insights that may be difficult to obtain through traditional gene-based methods. For instance, in the original publication of the melanoma dataset, the authors identified two clusters of malignant cells named “patient_specific_A” and “patient_specific_B” but could not infer their functional characteristics based on marker genes alone. By leveraging SeuratExtend’s pathway-based approach, we can easily identify the specific pathways that distinguish these clusters (**Figure 5C**), thereby gaining a deeper understanding of their biological roles.

**Figure 5.**
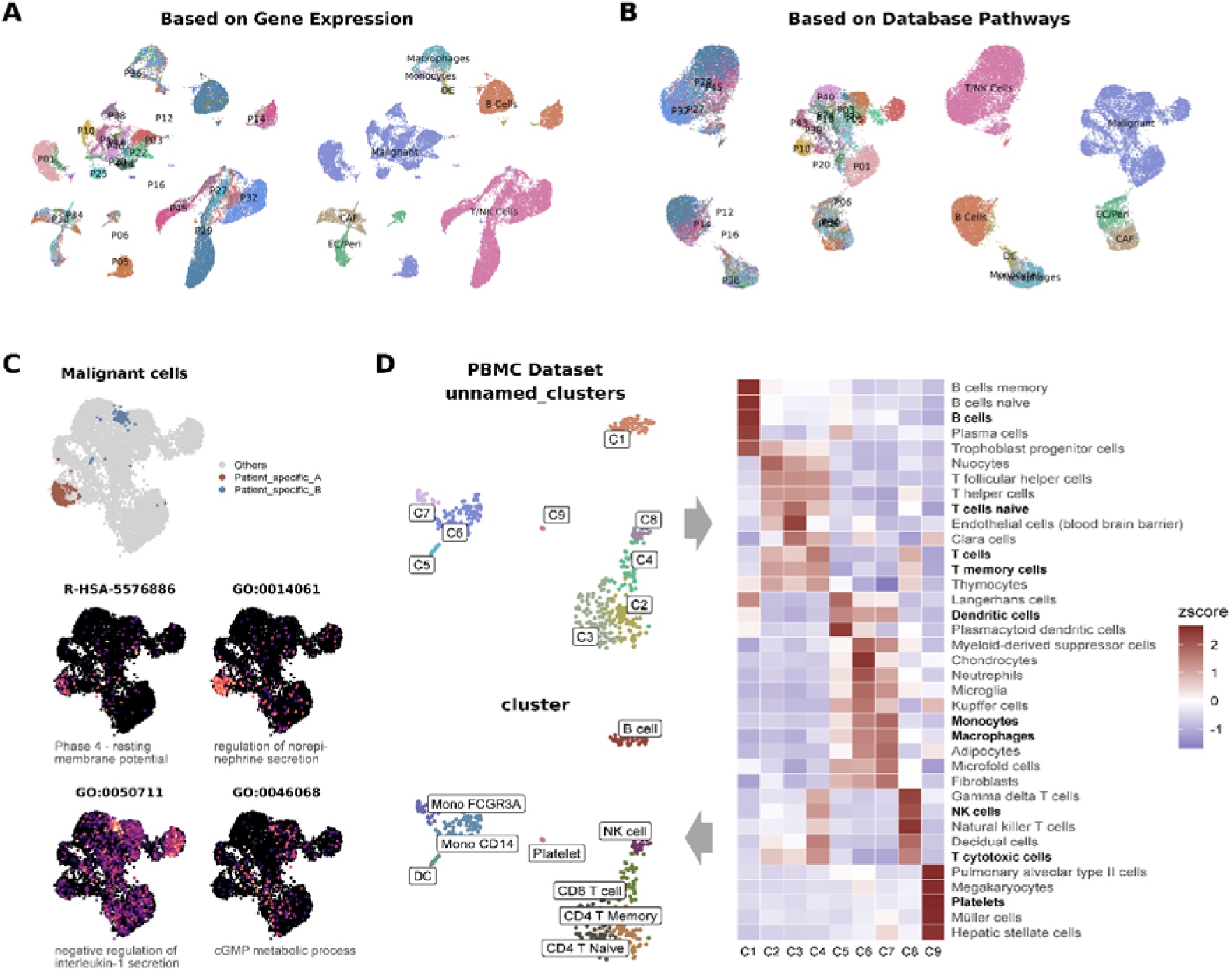
SeuratExtend’s novel applications in pathway-level analysis and cluster annotation. (A) UMAP visualization of melanoma scRNA-seq data colored by sample origin and clusters, demonstrating the influence of batch effects on gene expression-based clustering. (B) UMAP visualization of the same data colored by sample origin and clusters derived from pathway enrichment scores based on multiple databases (GO, Reactome, KEGG, and BioCarta), showing the grouping of cells with conserved features (e.g., T cells and B cells) while preserving biological variations in malignant cells. (C) UMAP visualization of malignant cells highlighting the location of patient_specific_A and patient_specific_B clusters (top) and their corresponding specific pathway activities (bottom). (D) UMAP visualization of unsupervised clustering results yielding nine unnamed clusters (top-left) and the corresponding heatmap of top enriched cell type signatures from PanglaoDB for each cluster (right), facilitating semi-automated cluster annotation (bottom-left).

SeuratExtend’s pathway-based approach is highly versatile, as it can be applied to various databases, as well as customized pathway databases, enabling fine-tuned clustering for specific datasets. However, it should be acknowledged that this method shares some of AUCell’s limitations, such as sensitivity to the number of genes expressed per cell (nGene). Consequently, substantial differences in nGene across samples may impact the computational results.

### Semi-Automated Cluster Annotation with Signature Enrichment Analysis

SeuratExtend also facilitates semi-automated cluster annotation without the need for additional tools like SingleR (Dvir Aran, Aaron Lun, Daniel Bunis, Jared Andrews, Friederike Dündar 2019). By leveraging databases such as PanglaoDB (Franzén, Gan, and Björkegren 2019), which contains marker genes for over 100 cell types, researchers can calculate AUCell scores to identify enriched cell type signatures within each cluster, greatly assisting the annotation process.

In the provided example, unsupervised clustering yields nine unnamed clusters (**Figure 5D**). By calculating AUCell scores using PanglaoDB and sorting the top cell type signatures for each cluster, researchers can efficiently annotate the clusters based on this information (**Figure 5D**).

In summary, SeuratExtend’s novel applications in pathway-level analysis and cluster annotation demonstrate its potential to address critical challenges in scRNA-seq data interpretation. By leveraging integrated pathway databases and signature enrichment analysis, SeuratExtend enables researchers to gain deeper insights into the functional characteristics of cell populations, streamline the annotation process, and ultimately expand the analytical horizons of single-cell transcriptomics.

## DISCUSSION

SeuratExtend represents a significant advancement in the field of single-cell RNA sequencing (scRNA-seq) analysis, addressing the critical challenges that have emerged from the rapid proliferation of computational tools and algorithms in this domain. Our comprehensive evaluation and integration of essential analytical components have resulted in a robust, user-friendly framework that streamlines complex workflows and enhances the accessibility of advanced scRNA-seq analysis techniques.

The development of SeuratExtend was guided by three core principles: integration, intuitive design, and visual aesthetics. By seamlessly incorporating multiple databases, analytical tools, and visualization techniques, we have created a cohesive ecosystem that bridges the gap between R and Python environments. This integration not only simplifies the analytical process but also expands the repertoire of available tools for researchers working primarily in R. The intuitive design of SeuratExtend, featuring straightforward functions and extensive documentation, significantly lowers the entry barrier for both novice and experienced users. Furthermore, our emphasis on visual aesthetics, exemplified by the carefully curated color schemes and optimized visualization methods, enhances the clarity and impact of data representation in scientific communications.

While SeuratExtend has integrated various essential components, including denoising, batch integration, pathway and functional analysis, gene regulatory network inference, trajectory and pseudotime analysis, it is not an all-encompassing encyclopedia-like toolbox. It does not currently cover all aspects of single-cell analysis, such as doublet removal, cell-cell communication, copy number variation inference, or automatic cluster annotation/label transfer. However, these features may be incorporated in future versions as SeuratExtend continues to evolve. The current version of SeuratExtend focuses on the most practical analyses distilled from real-world experience, complementing and optimizing existing tools, and crucially, establishing foundational frameworks like database integration and bridging the R and Python ecosystems. These robust foundations provide excellent scalability, positioning SeuratExtend with the potential to evolve into a comprehensive ecosystem surrounding Seurat, enabling users to perform end-to-end analyses while maintaining an intuitive and user-friendly experience.

To realize this potential, SeuratExtend’s future development and expansion will be greatly enhanced by active community engagement. As with many successful open-source projects, user feedback and contributions from the scientific community will play a crucial role in shaping the package’s trajectory. This collaborative approach ensures that SeuratExtend continues to evolve in alignment with users’ needs and the rapidly advancing field of single-cell genomics. We encourage users to provide feedback, suggest new features, and contribute to the codebase, fostering a vibrant ecosystem that can adapt to emerging challenges and opportunities in scRNA-seq analysis.

Looking ahead, SeuratExtend’s integration and intuitive nature position it as an excellent educational resource, further lowering the entry barrier for those eager to learn single-cell analysis. Additionally, the rise of large language models (LLMs) presents an opportunity for AI-assisted education. While SeuratExtend currently uses OpenAI’s platform for a chatbot, future endeavors may involve building more versatile chatbots using frameworks like Langchain and other LLMs like Claude and Llama 3, increasing accessibility and reducing costs. Furthermore, SeuratExtend’s standardized data analysis and visualization framework could pave the way for visual applications like Shiny apps, making scRNA-seq analysis accessible even to non-bioinformaticians.

In conclusion, SeuratExtend represents a significant stride in streamlining scRNA-seq data analysis, offering a comprehensive and integrated solution built upon the Seurat framework. By addressing the challenges of tool proliferation and complexity, SeuratExtend has made advanced scRNA-seq analysis more accessible to a broader range of researchers. With its robust foundations, scalable architecture, and commitment to user-friendliness, SeuratExtend is predestined to evolve into a comprehensive ecosystem, empowering researchers to harness the full potential of single-cell transcriptomics across various biological disciplines.

## DATA AVAILABILITY

The SeuratExtend package is freely available in the GitHub repository (https://github.com/huayc09/SeuratExtend) and has been deposited to figshare with the DOI: 10.6084/m9.figshare.26264255. The repository also contains comprehensive documentation and tutorials (https://huayc09.github.io/SeuratExtend/) to help users get started with the package and understand its functionalities. The example datasets used in the tutorials are available on Zenodo: https://zenodo.org/records/10944066.

## AUTHOR CONTRIBUTIONS

Y.H. conceived the idea, designed the study, developed the SeuratExtend package, performed data analysis, and drafted the manuscript. F.R. supervised the project and provided guidance throughout its development. L.W. provided extensive feedback on the package and reported bugs, contributing to its overall improvement. F.Z. assisted in testing the code and tutorials. All authors read and approved the final manuscript.

## ACKNOWLEDGEMENTS

Y.H. would like to express his sincere gratitude to the VIB KU-Leuven for their single-cell RNA-seq course, which provided the foundational knowledge essential for the development of SeuratExtend. Special thanks go to the individuals from the University Hospital Essen, KU Leuven-VIB Center for Cancer Biology, Institut National de la Santé et de la Recherche Médicale (Inserm), and Max Delbrück Center for Molecular Medicine (MDC) who contributed their time and effort to test the package. The author also acknowledges the assistance of Claude and ChatGPT in refining the code and manuscript language.

